# Comparison of changes in electrical activity, in isolated and *in vivo* hearts, induced by voluntary exercise in female rats

**DOI:** 10.1101/2020.11.16.385021

**Authors:** Rachel Stones, Mark Drinkhill, Ed White

## Abstract

Regular mild exercise is recommended to the general population as beneficial to health. Regular exercise typically leads to structural and electrical remodelling of the heart but in human studies it is difficult to relate the extrinsic and intrinsic influences on intact hearts to changes seen at the single cell level. In this study we wished to test whether changes in electrical activity in intact hearts, in response to voluntary wheel running exercise training, were consistent with our previous observations in single cardiac myocytes and whether these changes resulted in altered susceptibility to arrhythmic stimuli.

Female rats performed 5 weeks of voluntary wheel running. Implanted telemetry transmitters were used to measure electrocardiograms (ECGs) and determine heart rate variability (HRV) in conscious, unrestrained, trained (TRN) and sedentary (SED) animals. In isolated hearts, left ventricular epicardial monophasic action potentials (MAPs) were recorded and the responses to potentially arrhythmic interventions were assessed.

Exercise training caused cardiac hypertrophy, as indexed by a significantly greater heart weight to body weight ratio. Consistent with previous measurements of action potential duration in single myocytes, MAPs were significantly longer at 50%, 75% and 90% repolarization. Arrhythmic susceptibility was not different between SED and TRN hearts. Trained animals displayed significantly altered HRV by week 5, in a manner consistent with reduced sympathetic tone, however resting ECG parameters, including those most associated with repolarisation duration, were unaltered. We conclude that intrinsic changes to cellular cardiac electrophysiology, induced by mild voluntary exercise, are not attenuated by the electronic loading that occurs in intact hearts. However, *in vivo,* extrinsic neuro-hormonal control of the heart may minimize the effects of intrinsic alterations in electrical activity.

## Introduction

The numerous health benefits of regular, mild exercise has led to its recommendation to the general population (Fletcher *et al* 1996; Thompson *et al* 2003) and the associated changes in cardiac structure and electrical activity are usually seen as benign (Pelliccia *et al* 2000). However, some changes, such as hypertrophy and action potential prolongation can be pro-arrhythmic when associated with certain pathologies, such as hypertrophic cardiomyopathy, and implicated in sudden cardiac death in athletes (Maron & Pelliccia 2006). Furthermore, mechanisms associated with important exercise-induced changes in cardiac electrical activity are still disputed, for example, exercise-induced bradycardia is reported to result from altered autonomic balance (Billman 2009), but alternatively, from downregulation of HCN4 ion channels (Da Souza *et al* 2014). Therefore, better understanding of the electrical changes in cardiac activity in response to exercise training is still needed.

Most human and other mammalian studies of exercise have concentrated upon the effects of intense training regimes on the cardiovascular system (Moore & Korzick, 1995; Maron & Pellicci, 2006; Kemi *et al* 2008), with less study on the effects of mild exercise. The voluntary wheel running rat model is a well-established model of mild exercise (Scislo *et al* 1993; Rodnick *et al* 1998; DiCarlo *et al* 2002; Stones et al 2008b; Stones *et al* 2009; Natali *et al*, 2015; Wang & Fitts, 2017) but training–induced changes in cardiac electrical activity in this model are relatively less studied. Interestingly, given the situation in humans described above, we have previously reported cardiac hypertrophy and action potential prolongation of single myocytes isolated from the sub-epicardial myocardium of this model (Natali *et al* 2002; Stones *et al* 2009). These changes possibly occurring as a result of downregulation of the transient outward K^+^ current, I_to_ (Stones *et al* 2009).

However, given the structural and electrical complexity of the intact heart, compared to single myocytes, (Taggart *et al* 2003; Stones *et al* 2008a) it is important to verify that phenomena observed in isolated single myocytes persist in the intact heart. It is also important to note that in addition to any intrinsic changes that occur in the myocardium, the response of the heart to regular exercise is also modulated *in vivo,* by neuro-hormonal factors (Danson *et al* 2005). These extrinsic factors may amplify or attenuate intrinsic modulation, the separation of these factors is difficult to study in humans.

In this study we wished to characterise the changes in the APD and ECG of intact, isolated and *in vivo* hearts in the female voluntary running rat model. The purpose was to (i) compare with existing data from single cardiac myocytes and (ii) to test the hypothesis that in response to mild, voluntary exercise, intrinsic electrical remodelling in the intact rat heart persist *in vivo* in the presence of extrinsic regulation and (iii) that such changes are benign, with respect to selected arrhythmic stimuli.

## Methods

The investigation was performed in accordance with the recommendations of the Directive 2010/63/EU of the European Parliament on the protection of animals used for scientific purposes and local ethical approval.

### Exercise training model

Female Sprague-Dawley rats, approximately six weeks old, were weight and age matched, and randomly assigned to either a sedentary (SED) or trained (TRN) group. Female rats were used as they have been reported more likely to run spontaneously than male rats e.g. (Rodnick *et al* 1989; Wang & Fitts, 2017). TRN rats had free access to running wheels over a 5 week period as previously described (Natali *et al* 2002; Stones *et al* 2008b). Individual running distances were recorded daily. All animals were housed at 21°C with a 12:12hr light: dark cycle and had access to standard rat chow and water *ad libitum*.

### Telemetric recording of ECG

Surgical implantation of telemetry devices (TA10ETA-F20, Data Sciences International, St Paul, MN, USA) was conducted under isoflurane (2-3%) anaesthesia, using aseptic technique, 7-10 days prior to the training period. We implanted devices designed for use in mice, which are smaller and lighter than standard rat devices, in order to minimize impact on running behaviour (see Fig 1A). Electrocardiograms (ECGs) and activity were recorded in conscious, unrestrained TRN and SED rats for 2 hours per day for the duration of the training period. Data were acquired with Dataquest A.R.T. 4.1 Gold software (Data Sciences International).

**Figure 1.**
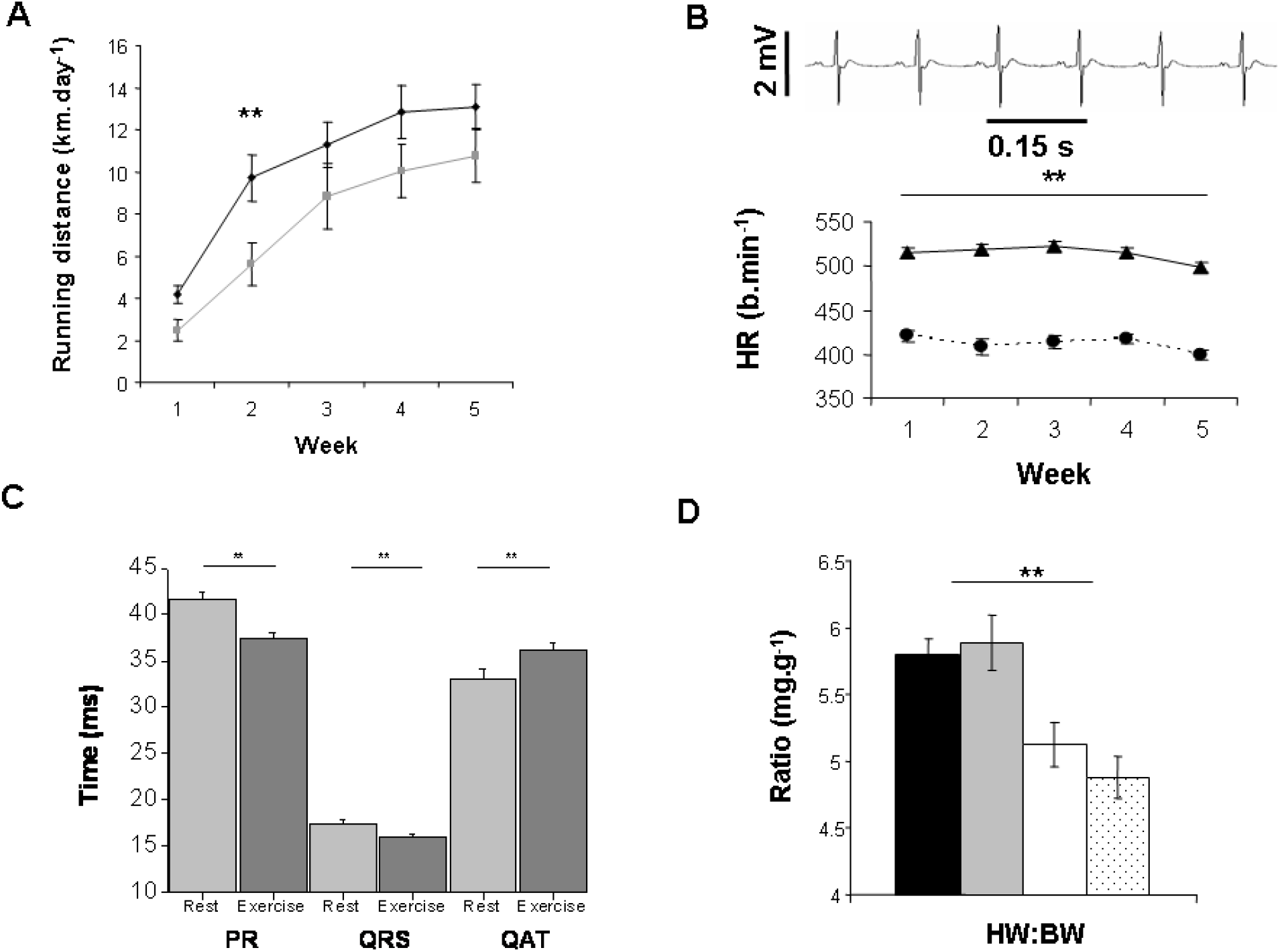
Running distances and ECG parameters for trained rats. Heart weight to body weight ratios (HW:BW) for trained and sedentary rats. Running distance increased gradually over the duration of the training period, for both non-implanted rats (♦, n=19) and rats implanted with telemetry devices (◼, n=12). Average daily running distance was only significantly different between these groups at week 2. (B) Upper panel shows a representative ECG recording in the II lead configuration recorded during rest from a conscious, unrestrained rat by radio telemetry. Lower panel compares heart rates at rest (⚫) and during exercise (▲) in TRN animals through the 5 week training period (**P < 0.001, n=12). (C) ECG parameters at rest and during exercise during week 5. The PR and QRS interval were significantly reduced during exercise whilst the QAT was significantly prolonged (** P = 0.001, n = 12). (D) Heart weight to body weight ratio (HW:BW) was significantly higher in both trained groups compared to both sedentary groups but was not significantly affected by implantation of telemetry devices (TRN non-implanted ◼, n=19; implanted ◻, n= 12; SED non-implanted ◻, n=17; implanted dotted, n=12, ** P <0.001).

ECG waveform data obtained during a resting period, as indicated by zero counts from the activity monitors incorporated in the ECG telemetry devices, was analysed with DSI Ponemah software (P3 Plus Version 4.4, Data Sciences International) to determine heart rate (HR, beats.min^−1^); PR (the time in ms from the start of the P wave to the start of the R wave); QRS (the duration in ms of the QRS complex); Q alpha T (QAT, time from the Q wave to the peak of the following T wave in ms and indicative of the shortest APD, typically the sub-epicardial APD); Ta (area of the T wave, ms.mV) and Tp (peak T wave amplitude, mV). Both Ta and Tp are indicative of transmural dispersion of APD (Yan & Antzelevitch, 1998). Tpe (time between the peak of the T wave to the end of the T wave in ms) was also measured, under the conditions of our experiment this parameter is thought to be indicative of the global dispersion in repolarization (Antzelevitch *et al* 2007; Optof *et al* 2007).

### Heart Rate Variability

Heart rate variability (HRV) was determined from 2 min. segments of ECG data collected at rest, 2 days per week, for 5 weeks (Chart 5 Pro software, ADInstruments, Colorado Springs, CO). Fast Fourier transformation was performed with Welch window with an FFT setting *n* = 1024 points, with 50% overlap. Spectral power was quantified within the frequency bands: total power 0 to 5 Hz; low frequency power (LF), 0.04 to 1 Hz; and high frequency power (HF), 1 to 3 Hz (Imai *et al* 2008). LF is thought to be indicative of sympathetic and parasympathetic tone whilst HF is indicative of parasympathetic tone (Stauss 2003).

### Monophasic action potential recordings in the whole heart

At the end of the training period, rats were weighed and killed by cervical dislocation. Hearts were removed and flushed with a bicarbonate-buffered solution of the following composition (mM): 118.5 NaCl, 14.5 Na HCO_3_, 4.2 KCl, 1.2 KH_2_PO_4_, 1.2 MgSO_4_7H_2_O, 11.1 Glucose, 1 CaCl_2_, pH 7.4, blotted dry and weighed before Langendorff perfusion at 0.11 ml.s^−1^.g^−1^ heart weight with the above solution at 37 °C.

A Franz-type monophasic action potential (MAP) electrode was used to record MAPs from the epicardial surface of the left ventricular wall at the mid-line using guidelines given by Franz (1999). In experiments where hearts were paced, a stimulation frequency of 5 Hz was applied via platinum electrodes close to the junction of the right atria and right ventricle. Pacing at slightly above intrinsic rates was used to negate any effect of heart rate upon MAP duration, to standardize the timing of rapid pacing with respect to prior MAPs and to monitor the generation of extra-systoles within a train of excitations (see below). Intrinsic heart rate and its variability were assessed by recording MAPs in unpaced hearts.

To assess arrhythmic susceptibility the response to two arrhythmic stimuli was measured. In paced hearts, a brief period of rapid pacing (2 s at 50 Hz) was interpolated into a train of stimulation at 5 Hz in order to disrupt rhythm e.g. (Ninio et al 2005; Noujaim et al 2007). In unpaced hearts ischemia/reperfusion was provoked by 30 min. of zero flow followed by reperfusion of the tissue (Carmeliet 1999). Our arrhythmic index was the standard deviation of the inter-beat interval (SD.IBI) calculated from the interval between MAP upstrokes (using Chart, ADInstruments) in the 10 s immediately prior to and following rapid pacing or immediately before and after implementation of the ischemia/reperfusion protocol. A perfectly rhythmic heart has a SD.IBI of zero because all upstroke to upstroke intervals are identical, deviation from this value, caused either by extra or dropped excitations, indicates a disruption to heart rhythm (Langer *et al* 1999). In paced experiments the number of extra-systoles (MAPs not triggered by the stimulator) was also recorded as a separate parameter.

### Statistical Analysis

All data are expressed as mean ± S.E.M unless otherwise stated. Statistical analysis was performed (with SigmaStat 3.5, Systat Software) by appropriate two way analysis of variance (ANOVA) or Student's paired or unpaired t-test or equivalent non-parametric tests if data were not normally distributed. Statistical significance was assumed with a probability of less than 0.05.

## Results

TRN animals ran voluntarily over the 5 week experimental period, reaching a peak daily distance of 13.01 ± 1.07 km.day^−1^ (Fig 1A). Average running speed was 14.1 ± 1.5 m.min^−1^ and heart rate increased by approximately 100 bpm during exercise throughout the 5 week period (see Fig. 1B). We observed that during exercise there was a significant shortening of both the PR interval and QRS complex but a significantly lengthened QAT, (P = 0.001, n =12) (Fig 1C), other measured parameters were not altered during exercise. These changes suggest faster atrial-ventricular conduction and ventricular excitation and a longer APD during wheel running exercise. Exercise training was associated with a decreased body weight in TRN animals (213 ± 3.3 g TRN; 224 ± 3.4 g SED, P < 0.05) but an increased heart weight (1.25 ± 0.03 g TRN; 1.13 ± 0.03 g SED, P <0.001 ANOVA, n = 31 TRN, n= 29 SED) hence there was cardiac hypertrophy as indexed by a significantly increased heart weight to body weight ratio in the TRN groups compared to SED groups (Fig 1D).

There was no significant difference in the level of wheel running (with the exception of distance run after 2 weeks) (Fig 1A) or the heart weight to body weight ratio between TRN animals implanted with telemetry devices and non-implanted TRN animals (Fig 1D). Our results indicate that implantation of the smaller mouse telemetry devices had minimal effects on the animals and that this is a viable technique for *in vivo* monitoring of unrestrained exercising rats.

The intrinsic beating frequency of excised hearts was not significantly different between TRN, 4.58 ± 0.14 Hz (n=11) and SED, 4.92 ± 0.2 Hz (n=11) hearts. We therefore paced hearts at slightly above intrinsic rates (5Hz) to negate any effect of heart rate upon MAP duration. Left ventricular epicardial MAP recordings were made from the mid-line (Fig 2A). MAP duration from TRN hearts was significantly longer than SED hearts at all levels of repolarization measured (Fig 2B) indicating there was intrinsic electrical remodelling of the intact heart in response to voluntary exercise.

**Figure 2.**
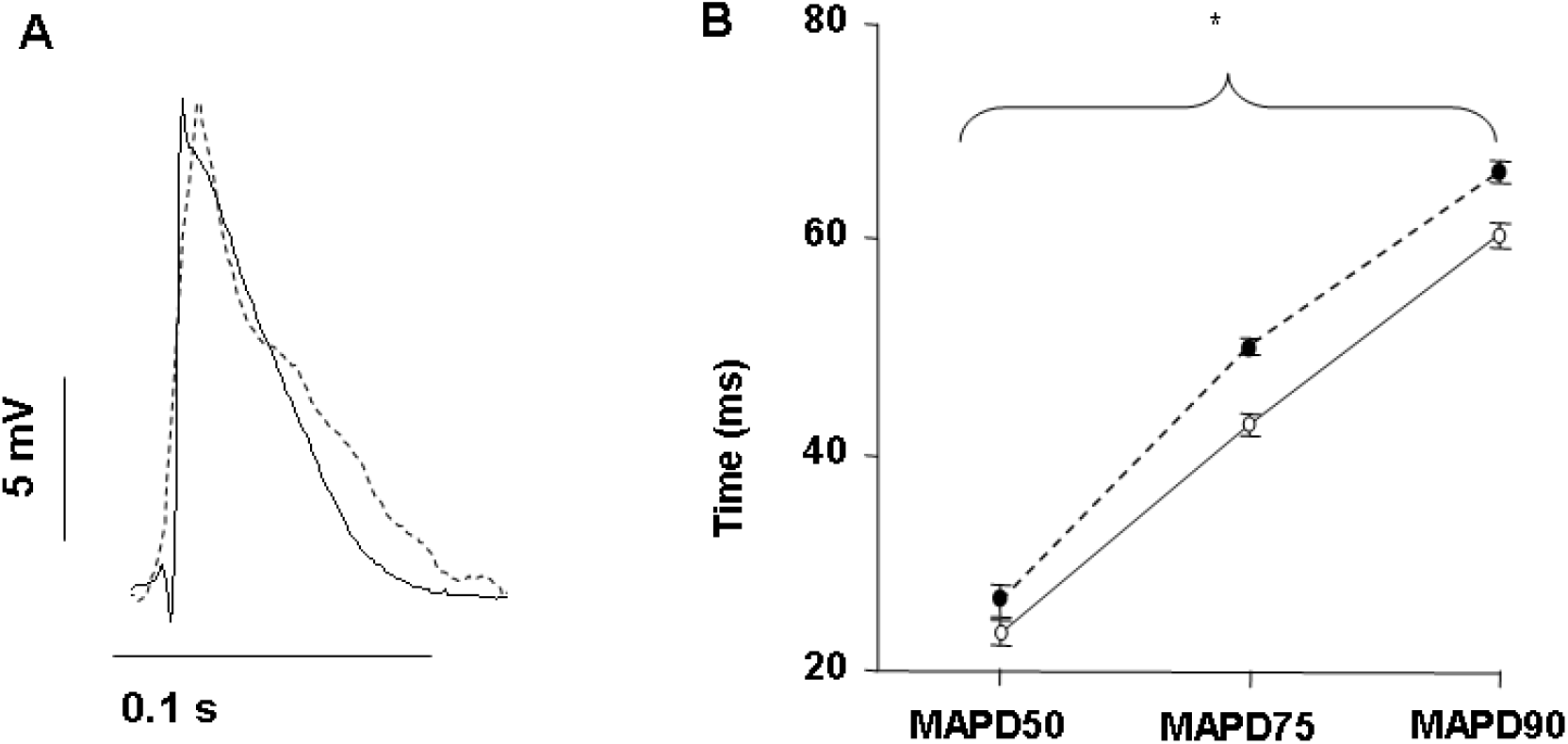
Monophasic action potential recordings from trained and sedentary rat hearts. (A) Monophasic action potentials (MAPs) were recorded from the left ventricular epicardial surface of trained (dotted line) and sedentary rat hearts (solid line) at 37°C at a stimulation frequency of 5 Hz. (B) MAPs recorded from the hearts of trained rats (●, n=12) were significantly longer than those measured from sedentary rat hearts (○, n=10) at 50, 75 and 90% repolarization (* P<0.05).

When rapid pacing (50 Hz for 2s) was applied in order to disrupt rhythm (Fig 3A) there was a significant increase in the SD.IBI, caused by both the generation of extra-systoles and by dropped excitations. However there was no significant difference between the post-rapid pacing SD. IBI of TRN and SED hearts (Fig 3B), nor in the number of extra-systoles generated (Fig 3C).

**Figure 3.**
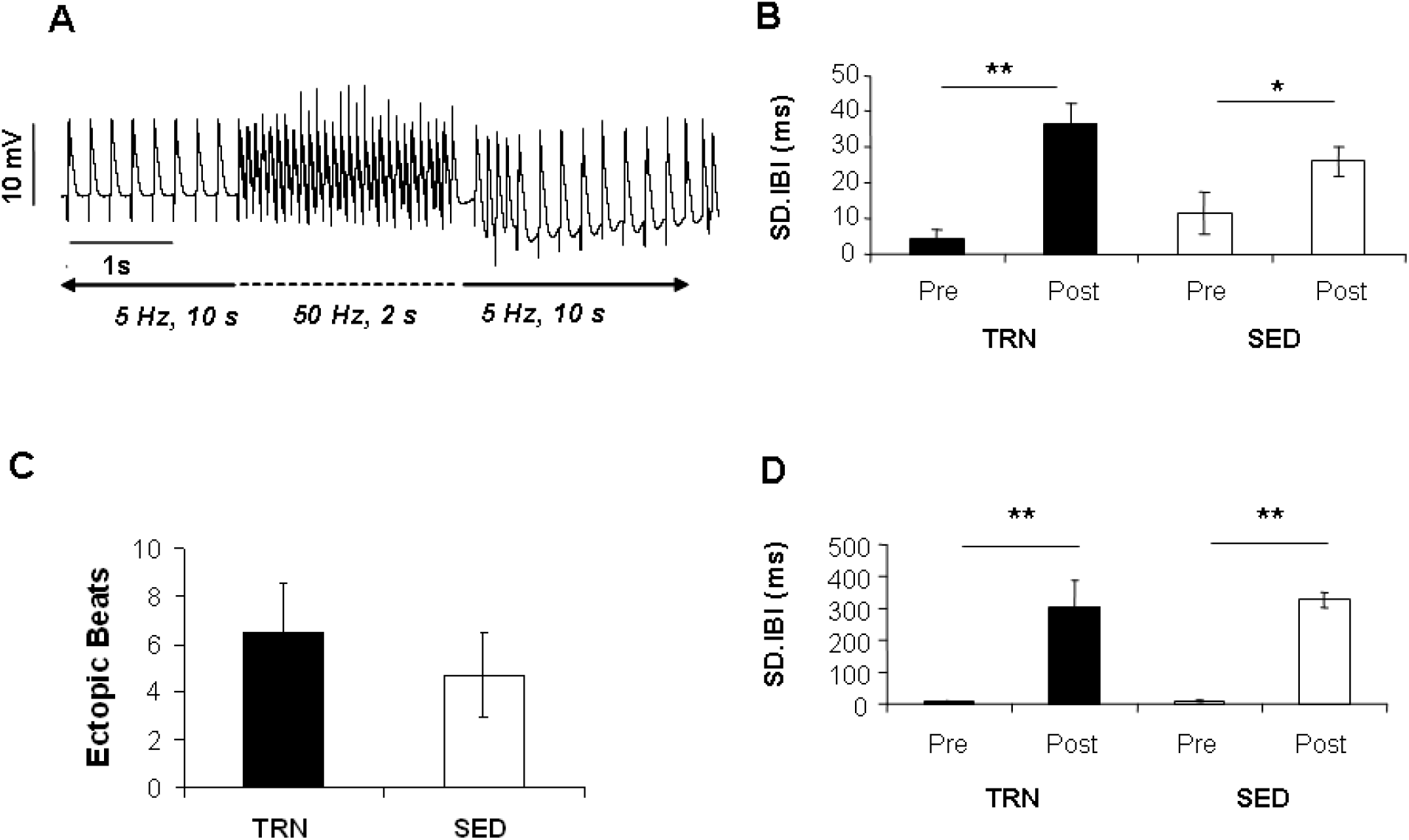
Arrhythmic susceptibility in trained and sedentary rat hearts. (A) Monophasic Action potential (MAP) recordings in an isolated heart showing that following the interpolation of a brief period of rapid stimulation (50 Hz for 2 s) there was a disruption to the rhythm of the heart. (B) The standard deviation of the inter-beat interval (SD.IBI) in the 10 s immediately following the period of rapid stimulation. (C) The number of ectopic beats occurring in the 10 s period immediately following rapid stimulation. Whilst SD. IBI and the number of ectopic beats was significantly increased in both trained (TRN) and sedentary (SED) rat hearts following rapid stimulation (*P < 0.05, **P < 0.001) there were no significant differences between TRN and SED responses (P > 0.05). (D) SD. IBI was recorded for 1 min. immediately prior (pre) to 30 min. ischemia and following reperfusion (post) in unpaced hearts. SD.IBI was small pre-ischemia/reperfusion and increased significantly post-ischemia/reperfusion. (**P < 0.001). However the pre- and post-responses did not differ between TRN and SED rat hearts (P > 0.05). Data from 9-11 SED and 9-11 TRN hearts.

In unpaced hearts the SD.IBI was small (indicating a regular sinus rhythm) and not significantly different between SED and TRN hearts (Fig 3D pre-). Application of the ischemia/reperfusion protocol lead to a substantial disruption of sinus rhythm and subsequent increase in SD. IBI but again, these responses did not differ between SED and TRN hearts (Fig 3D post-). Therefore despite the presence of both hypertrophy and prolonged electrical repolarisation, TRN hearts were neither pre-disposed, nor protected, from arrhythmia.

In addition to factors intrinsic to the heart, the cardiac response to exercise *in vivo* is modulated by extrinsic neuro-hormonal factors. HRV has components that are influenced by parasympathetic tone (HF component) and sympathetic/parasympathetic tone (LF component). We observed no changes in the HF component of HRV. However, a gradual rise in the LF component, that was seen in the SED animals, was suppressed in the TRN animals such that there was a statistically significant difference in the LF component of SED and TRN animals by week 5 (Fig 4). Because the voluntary exercise model does not allow comparison of ECGs between TRN and SED animals at equivalent levels of exercise, we compared their ECG profiles at rest. There were no significant changes in resting heart rate or ECG parameters between TRN and SED animals (Table 1 and see Discussion).

**Figure 4.**
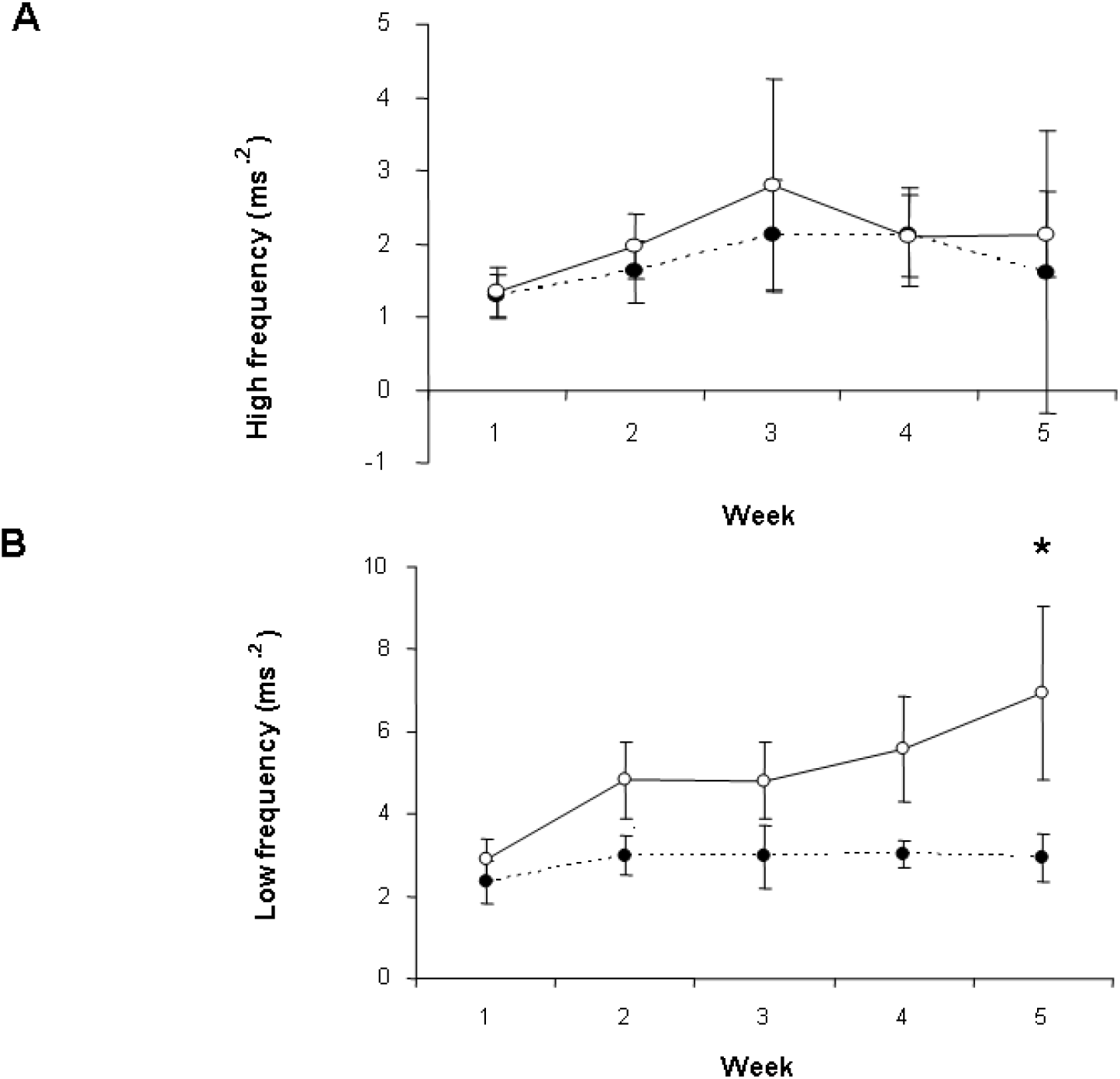
Heart Rate Variability in trained and sedentary rats. Heart rate variability (HRV) was recorded at rest from trained (●) and sedentary rats (○) and spectral power analysis performed (low frequency (LF) power, 0.04 to 1 Hz; and high frequency (HF) power, 1 to 3 Hz) to determine the influence of parasympathetic (HF) and sympathetic /parasympathetic (LF) tone respectively. Whilst there was no change in the high frequency component of HRV with training, there was divergence in the LF component that was statistically significant by week 5 (*P < 0.05, n=12).

**Table 1.** ECG parameters. ECGs were analyzed at rest (as indicated by zero activity from the activity monitors incorporated into the telemetry devices). We observed no changes in the resting heart rate or resting ECG parameters between TRN or SED animals (P < 0.05, n=12). The parameters measured were: heart rate (HR, beats.min^−1^); Q alpha T (QAT, time from the Q wave to the peak of the following T wave in ms and indicative of the shortest APD, which is typically the sub-epicardial APD); Ta (area of the T wave, ms. mV); Tp (peak amplitude of the T wave, mV) and Tpe (time between the peak of the T wave to the end of the T wave in ms). Tp, Ta and Tpe would be expected to decrease if there was a decrease in the transmural dispersion of repolarization in response to a lengthening of epicardial APD and Tpe altered by any changes in global dispersion of repolarization.

|  | HR (b.min<sup>-1</sup>) |  | QAT (ms) |  | Ta (ms.mV) |  | Tp (mV) |  | Tpe (ms) |  |
| --- | --- | --- | --- | --- | --- | --- | --- | --- | --- | --- |
| Week | SED | TRN | SED | TRN | SED | TRN | SED | TRN | SED | TRN |
| 1 | 426.4±6.6 | 421.2±6.6 | 32.5±1.0 | 32.3±0.9 | 2.2±0.3 | 2.3±0.3 | 0.04±0.031 | 0.056±0.033 | 19.1±1.3 | 18.5±0.7 |
| 2 | 420.3±8.0 | 408.7±8.7 | 31.6±1.0 | 34.4±1.7 | 1.9±0.1 | 2.3±0.3 | 0.032±0.033 | 0.026±0.043 | 17.3±0.9 | 20.0±1.4 |
| 3 | 394.7±9.1 | 414.3±7.5 | 33.7±1.6 | 32.5±1.0 | 1.6±0.1 | 2.5±0.3 | 0.035±0.023 | 0.02±0.04 | 16.9±1.0 | 19.1±1.5 |
| 4 | 397.3±8.6 | 418.1±6.2 | 35.2±1.9 | 33.5±0.8 | 1.7±0.2 | 2.2±0.3 | 0.006±0.028 | 0.024±0.035 | 17.7±0.9 | 19.5±1.0 |
| 5 | 389.6±8.4 | 399.8±5.8 | 33.7±1.3 | 33.0±1.1 | 1.7±0.2 | 2.2±0.3 | 0.006±0.026 | 0.007±0.036 | 17.0±0.9 | 18.1±1.0 |

## Discussion

Regular mild/moderate exercise is recommended to the general population by health care professionals (Fletcher *et al* 1996; Thompson *et al* 2003) therefore a full understanding of the response of the heart to mild exercise is important if its full benefits are to be reaped. The model of voluntary wheel running in rats is well established (Scislo *et al* 1993; Rodnick *et al* 1998; DiCarlo *et al* 2002; Stones et al 2008b; Stones *et al*, 2009, Natali *et al*, 2015, Wang & Fitts, 2017). It is designed to investigate the effects of mild, non-enforced exercise as it has been shown that the responses of rodents to voluntary exercise can differ from those caused by enforced exercise e.g. (Yancey & Overton 1993; Moraska *et al* 2000; Li *et al* 2014).

We observed that the use of mouse telemetry devices, in rats, did not significantly reduce either wheel running or the exercise-induced cardiac hypertrophy. This approach also allowed us to report ECG parameters from this model. The increase in QAT suggests a lengthening of action potential during exercise consistent with some single cell and modelling studies on the relationship between action potential frequency and duration (see Salle *et al* 2008). Although at high stimulation frequencies (10 Hz) ADP has been reported to fall possibly as a result of increased IKATP (Wang & Fitts, 2017).

Left ventricular epicardial MAP duration was prolonged in the intact, isolated, trained heart. When compared to responses observed from isolated single myocytes, responses in whole hearts can be attenuated, by the electronic loading of cell neighbours in multicellular preparations (Taggart *et al*, 2003). However, in this case, the prolongation of MAPs from whole hearts compliments the prolonged action potential recorded from single cardiac myocytes (Natali *et al* 2002; Stones *et al*, 2009) indicating that intrinsic electrical re-modelling persists in the intact heart. It seems unlikely that these changes in electrical activity are caused by altered [Ca^2+^]i handling as we have reported key elements of [Ca^2+^]i handling unaltered in this model (Stones *et al* 2008b), rather we have shown a downregulation in I_to_ (Stones et al, 2009). This current is implicated in many models of rat cardiac hypertrophy (Bryant *et al* 1999; Wagner *et al* 2007). Studies using intense exercise models have reported alterations in outward K^+^ currents (Jew *et al* 2001) with no change in inward Ca^2+^ current (Mokelke *et al* 1997).

Rat models that induce hypertrophy and action potential prolongation via factors associated with disease, such as hypertension e.g. (Brooksby *et al* 1993; Shipsey *et al* 1997; McCrossan et al, 2004) are known to be more susceptible to arrhythmogenic stimuli even in the compensated stage (Evans *et al* 1995). In contrast, the combination of hypertrophy and MAP prolongation induced by voluntary exercise did not make the TRN hearts more (or less) susceptible to the arrhythmogenic stimuli. The lack of protection seen in our study contrasts with that of (Beig *et al* 2011) where voluntary exercise in male rats improved resistance to arrhythmias induced by the Na^+^ channel opener aconitine. Enforced exercise regimes have also shown improved responses to arrhythmic stimuli in both normal rats (Dor-Haim *et al* 2013) and rats with myocardial infarction (Dor-Haim *et al* 2017) and in the Zucker Diabetic Fatty rat with a running distance of only 3km per week (Van Hoose *et al* 2010). It is worthy of note that in our study the mean running distance was over 90 km per week.

The changes we have observed may be related to the benign alterations in cardiac electrical activity seen in some human subjects (Pelliccia *et al* 2000) and suggest the voluntary wheel running model is a useful tool for the investigation of mild responses to exercise. Hypertrophy caused by physiological and pathological stimuli have distinct signalling pathways and outcomes (McMullen & Jennings 2007). Our data are consistent with an important distinction between the beneficial response to voluntary exercise and to the previously reported responses to pathological causes of hypertrophy and electrical remodelling e.g. (Evans et al 1995).

We observed no differences between the resting HR of TRN and SED animals *in vivo*. Although resting heart rate bradycardia is often cited as a characteristic of the cardiovascular response to exercise training (Collins & DiCarlo; 1997; Chandler & DiCarlo 1998) it should be noted that this change is not always observed (Negrao *et al* 1993) especially in females (Chen & DiCarlo 1996) suggesting an influence of gender in the response to exercise. Despite an exercise-induced alteration in epicardial APD recorded in intact, excised hearts at physiological temperatures and stimulating frequencies, there was no manifestation of this change in ECG parameters recorded in conscious, unrestrained animals. The parameter most closely linked to epicardial APD (QAT) was not significantly altered. Additionally there was no evidence for an alteration in the transmural or global dispersion of repolarization (which might be expected if epicardial APD changed) because Tp, Ta and Tpe were not altered (Yan & Antzelevitch 1998; Antzelevitch *et al* 2007).

Our observations therefore suggest that *in vivo,* neuro-hormonal influences upon the heart attenuate intrinsic changes induced by mild exercise. Our investigation of HRV provided no evidence for a change in parasympathetic tone, consistent with a lack of change in resting heart rate but showed a significant suppression of the rise in the LF component of HRV. This is indicative of a decrease in sympathetic tone in response to voluntary exercise training and is consistent with our reported decrease in sensitivity to β2-adrenoceptor stimulation in single ventricular myocytes in this training model (Stones et al 2008b). Our observation of maintained parasympathetic tone or HF power with depressed sympathetic tone or LF power is also consistent with previous reports (Scislo et al 1993; Collins & DiCarlo 1997; DiCarlo et al 2002). In conclusion, we found that in intact, isolated whole hearts at physiological temperature and stimulation frequency, MAP duration was lengthened in response to mild exercise training, however, these intrinsic changes in electrical activity may be masked by extrinsic factors *in vivo.* The ability to investigate intrinsic cellular and molecular mechanisms of the response to mild exercise in human cardiac tissue is limited, therefore animal models also displaying a mild response to exercise, such as the voluntary wheel running rat model, provide useful information in this important field of study.

## Acknowledgements

This work was supported by The Wellcome Trust

